# NONU-1 encodes a conserved endonuclease required for mRNA translation surveillance

**DOI:** 10.1101/674358

**Authors:** Marissa L. Glover, A. Max. Burroughs, Thea A. Egelhofer, Makena N. Pule, L. Aravind, Joshua A. Arribere

**Author notes:** Corresponding Author **Contact Information**: Joshua A. Arribere Sinsheimer Labs, University of California at Santa Cruz 1156 High St, Santa Cruz, CA 95064.

## Abstract

Cellular translation surveillance rescues ribosomes that stall on problematic mRNAs. During translation surveillance, endonucleolytic cleavage of the problematic mRNA is a critical step in rescuing stalled ribosomes. However, the nuclease(s) responsible remain unknown. Here we identify NONU-1 as a novel endoribonuclease required for translation surveillance pathways including No-Go and Nonstop mRNA Decay. We show that: (1) NONU-1 reduces Nonstop and No-Go mRNA levels; (2) NONU-1 contains an Smr RNase domain required for mRNA decay and with properties similar to the unknown endonuclease; and (3) the domain architecture and catalytic residues of NONU-1 are conserved throughout metazoans and eukaryotes, respectively. We extend our results in *C. elegans* to homologous factors in *S. cerevisiae*, showing conservation of function of the NONU-1 protein across billions of years of evolution. Our work establishes the identity of a previously unknown factor critical to translation surveillance and will inform mechanistic studies at the intersection of translation and mRNA decay.

## INTRODUCTION

Numerous mechanisms exist to protect cells from the negative effects of errors in gene expression. Among these are translation surveillance pathways in which a ribosome identifies an early stop codon (Nonsense-Mediated mRNA Decay, NMD), a lack of stop codons (Nonstop Decay), or a block during translation elongation (No-Go Decay). Central to both Nonstop and No-Go Decay is the process of ribosome stalling. Recent work has also shown that ribosomes stall during Nonsense-Mediated mRNA Decay, effectively funneling NMD targets into Nonstop Decay (Hashimoto et al. 2017; Arribere and Fire 2018). Despite substantial mechanistic insight into translation surveillance pathways (reviewed in (Joazeiro 2017)), how ribosomal stalling communicates with mRNA decay machinery remains a central unsolved question.

Mounting evidence points to endonucleolytic cleavage of the mRNA in the vicinity of stalled ribosomes as an important early event in translation surveillance (*e.g.*, (Doma and Parker 2006; Guydosh and Green 2014; Ikeuchi et al. 2019; Navickas et al.)). While the identity of the critical nuclease(s) remain unknown, target mRNAs are eventually cleared in part by 3’>5’ degradation facilitated by the SKI RNA helicase in conjunction with the exosome (Doma and Parker 2006; van Hoof et al. 2002; Hashimoto et al. 2017; Arribere and Fire 2018). Knowledge of the identity and functions of factors that interface between translation and mRNA decay will illuminate a critical junction in gene expression and regulation. Identification of nuclease(s) at this junction would therefore significantly advance our understanding of translation, surveillance, and how targeted mRNA degradation occurs.

Here we identify a mutation that blocks Nonstop and No-Go mRNA Decay in *C. elegans*. The mutation identifies a new gene *nonu-1* whose structure predicts that it encodes a conserved nuclease component of translation surveillance. Furthermore, we show that the previously uncharacterized *S. cerevisiae* homologs *CUE2* and *YPL199C* contribute to Nonstop Decay in that organism. NONU-1 contains a conserved IF3-C fold domain previously implicated in processing RNA. Our results identify a critical new component of the translation surveillance machinery in two model organisms and suggest why this factor has been recalcitrant to discovery in *S. cerevisiae*.

## RESULTS

### *nonu-1* encodes a novel factor required for Nonstop mRNA Decay

We previously developed a phenotypic reporter in *C. elegans* that allowed us to identify Nonstop mRNA Decay factors via reverse and forward genetics (Figure 1A, (Arribere and Fire 2018)). Briefly, the reporter was constructed using the *unc-54* locus as expression and function of this gene has been extensively studied (Brenner 1974; Epstein et al. 1974; Dibb et al. 1985, 1989; Moerman et al. 1982; Bejsovec and Anderson 1988; Anderson and Brenner 1984) and *unc-54* has been used in previous suppressor screens (Hodgkin et al. 1989). To construct a Nonstop reporter at the *unc-54* locus we first integrated a GFP at the C-terminus of UNC-54 by CRISPR/Cas9. We also removed all stop codons from the 3’UTR and integrated a ribosomal skipping T2A sequence between the C-terminal GFP and the remainder of the 3’UTR. The T2A sequence is a viral-derived peptide that cotranslationally releases the upstream protein and allows UNC-54::GFP to escape Nonstop protein decay (so-called “Ribosome Quality Control”, (Bengtson and Joazeiro 2010; Shao et al. 2013; Shen et al. 2015)). We hereafter refer to the *unc-54::gfp::t2a::nonstop* reporter as *unc-54(Nonstop)*. Animals with the *unc-54(Nonstop)* reporter deficient in Nonstop mRNA Decay exhibit derepression of the locus, as evidenced by increased GFP fluorescence, mRNA expression, and egg laying (*unc-54* encodes a muscle myosin required in the vulva for egg laying) (Figure 1B, (Arribere and Fire 2018)).

**Figure 1:**
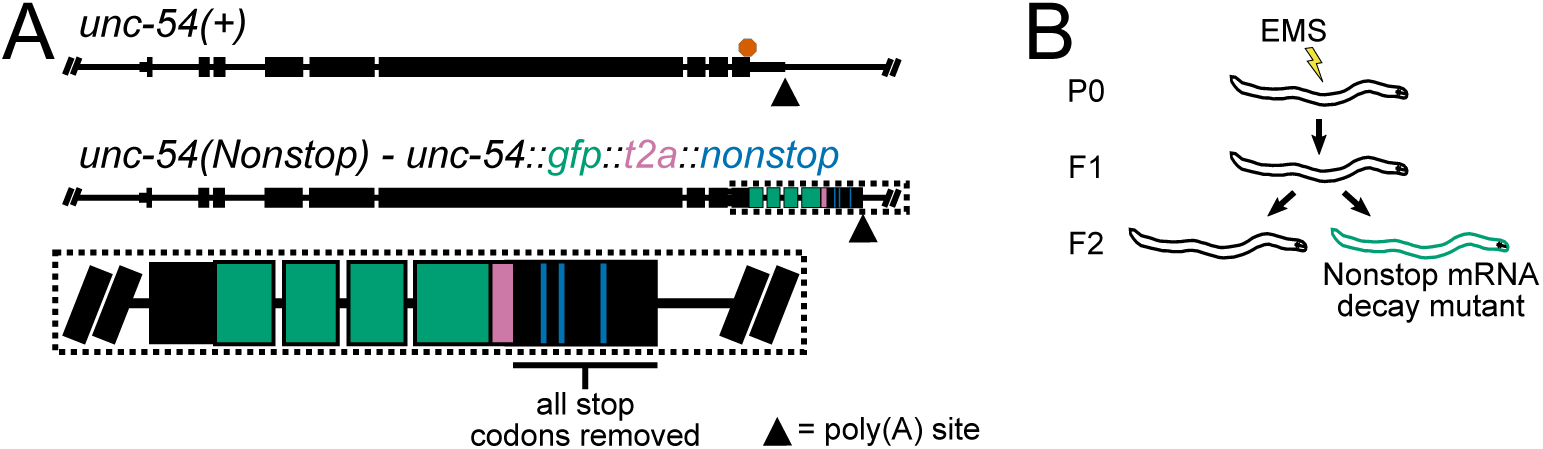
Nonstop mRNA Decay reporter and genetic screen. A: Gene diagrams showing annotated exons (black rectangles) at the wild type *unc-54* locus (top) and the *unc-54(Nonstop)* reporter (bottom). Red octagon indicates the location of the stop codon. Triangles indicate poly(A) sites. Inset zooms in on the region of interest, showing exons of GFP (green), T2A sequence (pink), and stop codons re-coded to sense codons (blue). B: Cartoon depicting the genetic screen to identify Nonstop mRNA Decay mutants in *C. elegans*.

Although our initial screen successfully identified *C. elegans*’ *skih-2* and *ttc-37* (homologs of *S. cerevisiae SKI2* and *SKI3*, respectively), it did not identify a factor that could function as an endonuclease. During a candidate approach leading to the identification of another translation surveillance factor *pelo-1* (homolog of Dom34/Pelota, (Arribere and Fire 2018)), we discovered that the dynamic range of GFP derepression in the *unc-54(Nonstop)* reporter for authentic surveillance mutants was greater than the range of our original screen. With this knowledge, we increased our ability to detect lower levels of GFP in order to detect derepression of the reporter similar to that observed in *pelo-1* animals.

We repeated the genetic screen and isolated an additional 36 mutants. We genetically mapped the causative locus in each strain by backcrossing to a polymorphic strain (also called Hawaiian variant mapping, Figure S1, Materials and Methods, (Arribere and Fire 2018; Doitsidou et al. 2010)). The majority of mutants mapped to loci homologous to known Nonstop mRNA Decay factors in other systems. However, two strains (WJA0675 and WJA0641) harbored mutations that mapped to an area of chromosome three lacking obvious homologs to known Nonstop mRNA Decay components (Figure S1). Visual inspection revealed that strain WJA0675 contained a Trp>STOP mutation in predicted ORF *f26a1.13*, and strain WJA0641 contained a Trp>STOP mutation in the neighboring ORF *f26a1.14* (Figure S2A).

Our subsequent analyses showed that *f26a1.13* and *f26a1.14* are a single gene that is required for Nonstop mRNA Decay: (1) EST, RNA-seq, and Ribo-seq data support the existence and translation of an mRNA transcript that spans both loci (Figure S2, (Hendriks et al. 2014)); (2) the unannotated transcript encodes a single open reading frame conserved in other metazoans and choanoflagellates; (3) an in-frame premature stop codon (inserted by CRISPR/Cas9) between *f26a1.13* and *f26a1.14* exhibited the same fold derepression of the *unc-54(Nonstop)* reporter observed in either mutant found in the genetic screen (WJA0675 or WJA0641, Figure 2A,B,C, S2A); (4) during the course of this work, *C. elegans* genome curators independently re-annotated *f26a1.13* and *f26a1.14* as a single gene (*f26a1.13*) based on RNA-seq data (Lee et al. 2018). We therefore conclude that *f26a1.13* and *f26a1.14* encode a single gene required for Nonstop mRNA Decay, and we hereafter refer to this gene as *nonu-1* (*nonu*, NOnstop NUclease). Homology searches with the encoded NONU-1 protein identified homologous proteins in diverse eukaryotes, but no homolog known to function in Nonstop mRNA Decay.

**Figure 2:**
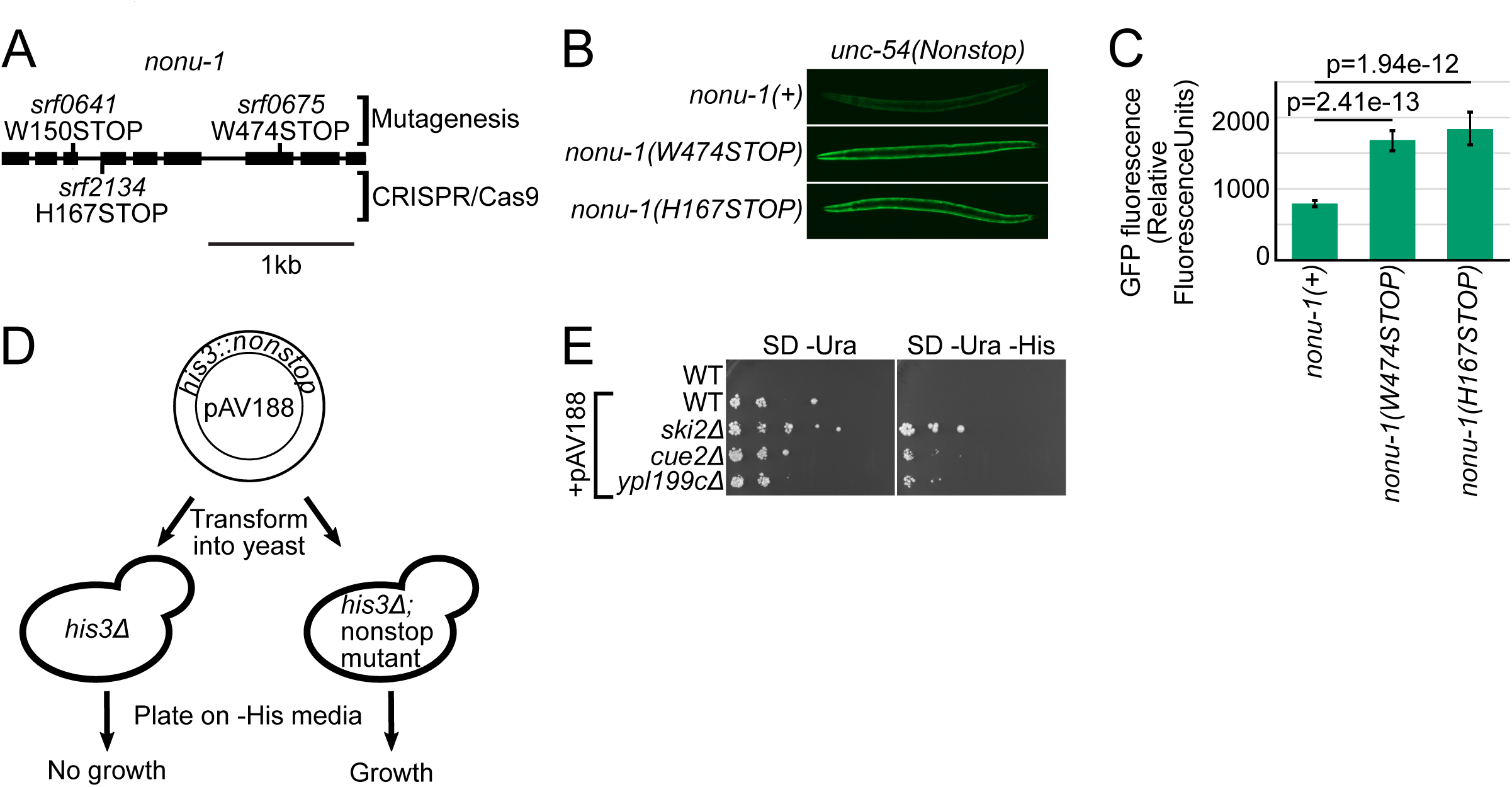
NONU-1 and its homologs in *S. cerevisiae* have a conserved function in Nonstop Decay. A. Gene diagram showing annotated exons (black rectangles) at the *nonu-1* locus. Mutations found via EMS mutagenesis are diagrammed above the gene, and a mutation generated by CRISPR/Cas9 is below the gene. Scale bar shows 1kb. B. Images of GFP expression in the indicated strains. See Methods. C. Quantification of GFP expression for strains shown in (B). Each bar represents an average of at least 5 images of 5 different animals, with 95% Confidence Intervals shown as error bars. P-values from Student’s T-test comparing mutants to *nonu-1(+)*. D. Diagram of the *S. cerevisiae* Nonstop Decay assay. *his3::nonstop* reporter plasmid (pAV188) that was transformed is shown on top. Yeast cartoons depict a *his3Δ* strain and a *his3Δ* strain that is also a Nonstop Decay mutant. Expected results of growing transformed stains on -His media are shown on the bottom. E. The indicated strains were transformed with the *his3::nonstop* reporter plasmid (pAV188) with *ski2Δ* as a positive control. Defects in Nonstop Decay manifest as improved growth on histidine dropout medium. Strains were serially diluted and plated on the indicated media. Pictures are representative of biological triplicates of the experiment.

### *S. cerevisiae* homologs of NONU-1 are required for Nonstop mRNA Decay

A previous genetic screen in *S. cerevisiae* failed to identify a homolog of NONU-1 (Wilson et al. 2007). We performed a homology search and identified two candidate homologs in *S. cerevisiae*: *YPL199C* and *CUE2*. To determine whether *CUE2* and/or *YPL199C* function in Nonstop mRNA Decay, we assayed the ability of mutant strains to derepress a *his3::nonstop* reporter previously used to identify and study factors required for Nonstop Decay in *S. cerevisiae* (Wilson et al. 2007; van Hoof et al. 2002). The *his3::nonstop* reporter encodes the *HIS3* gene with a recoded 3’UTR lacking any stop codons. In a strain competent for Nonstop Decay, the *his3::nonstop* reporter fails to complement a *his3Δ* strain and cannot grow without histidine. However, in a Nonstop Decay mutant, the *his3::nonstop* reporter is derepressed and can support growth of a *his3Δ* strain in the absence of histidine (Figure 2D).

Consistent with previous work, we observed substantial derepression of the reporter in a *ski2* mutant (Figure 2E). In either a *cue2Δ* or *ypl199cΔ* mutant we observed suppression of the *his3::nonstop* reporter. The magnitude of the suppression was significantly less than that conferred by a *ski2* mutation but was reproducible across independent isolates and technical replicates. The small magnitude of growth suppression compared to other mutants (*e.g.*, *ski2*) may have precluded either gene from being identified in a previous genetic screen (Wilson et al. 2007). We conclude that NONU-1 and its homologs in *S. cerevisiae* have a conserved function in Nonstop Decay. While *CUE2* and *YPL199C* each had a consistent effect on the *his3::nonstop* reporter, we note that the magnitude of this effect was below that of other factors (*i.e.*, *SKI2*), suggesting multiple independent mechanisms exist to repress Nonstop mRNAs.

### Domain architecture of NONU-1

To gain insight into NONU-1 function, we examined the domain structure of the protein and its metazoan orthologs and found that they contain several conserved domains (from N-terminus to C-terminus, Figure 3A, S3). The NONU-1 protein family is characterized by:

(1) An N-terminal basic region similar to a ribosome-binding motif at the N-terminus of the ribosomal protein S26AE. This basic stretch is only observed in the chordate versions of NONU-1 and is thus not pictured in Figure 3A. The basic stretch suggests that NONU-1 may bind directly to ribosomes.
(2) A domain of the P-loop kinase superfamily belonging to the polynucleotide kinase (PNK) clade. These kinase domains are known to phosphorylate RNA/DNA ends (Leipe et al. 2003; Burroughs and Aravind 2016). The P-loop kinase domain suggests NONU-1 may modify Nonstop mRNAs or their degradation products.
(3) Two ubiquitin-binding CUE (Coupling of Ubiquitin to ER degradation) domains of the UBA-like fold (Kang et al. 2003). Ub chains are an important signal for ribosomal stalling and suggest a mechanism of specificity for NONU-1 recruitment to stalled ribosomes (Ikeuchi et al. 2019; Simms et al. 2017; Garzia et al. 2017; Juszkiewicz et al. 2018; Saito et al. 2015).
(4) An Smr domain, homologous in structure and sequence to domains known to bind, cleave, or process RNA (Figure 3B, C, S3A, (Aravind et al. 2003)). Smr domains of some proteins function as an endoribonuclease (Bhandari et al. 2011; Zhou et al. 2017; Wu et al. 2016). (The NONU-1 Smr domain co-occurs with a highly charged, small helical extension that likely represents an extension of the Smr domain (“N-ext”, also known as “DUF1771”).) The existence of a domain known to function as an endoribonuclease makes NONU-1 a prime candidate for the unknown endonuclease involved in translation surveillance.

**Figure 3:**
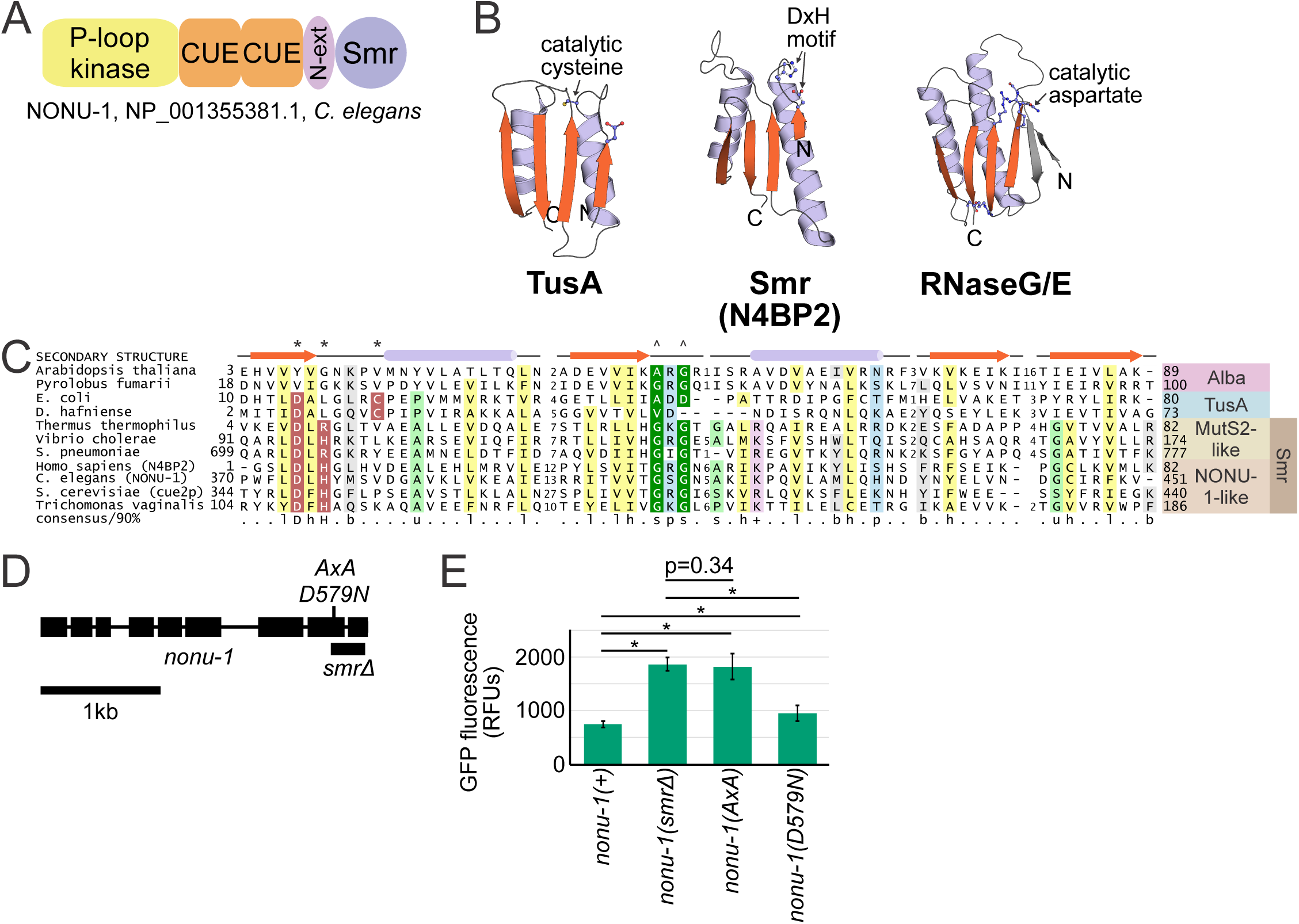
*nonu-1* contains a conserved Smr domain required for Nonstop mRNA Decay. A. Protein domain schematic of NONU-1. See text for details of each domain. B. Structures of representatives of the IF3-C fold domain, including a structure of Smr domain from the human homolog of NONU-1, N4BP2. Two of the structures shown are from domains known to contribute to RNA modification (TusA) or cleavage (RNaseE) with catalytic residues highlighted. The DxH motif of human N4BP2 is also highlighted. See Figure S3C for the phylogenetic relationship between these and other domains. C. Multiple sequence alignment depicting conserved motifs of the TusA-Alba-Smr assemblage of Smr domains. Protein secondary structure diagram is indicated above the sequence, and conserved amino acids are highlighted. * and ^ indicate amino acids known to be involved in catalysis and substrate recognition (respectively) in TusA or MutS2-like proteins. D. Gene diagram showing annotated exons (black rectangles) at the *nonu-1* locus. Mutations generated by CRISPR/Cas9 are shown (D579N above, *smrΔ* below). Scale bar shows 1kb. E. Quantification of GFP expression for the indicated strains (as in Figure 2C). P-values from Student’s T-test comparing mutants to *nonu-1(+)*. RFUs shown for Relative Fluorescence Units.

The combination of these domains characterizes the NONU-1 family of proteins found throughout metazoans and choanoflagellates as an endoribonuclease with a conserved role in diverse cell types and organisms.

### Catalysis by the Smr domain of NONU-1 is required for Nonstop mRNA Decay

Given the above observations, we investigated whether the Smr domain is important for NONU-1 function in Nonstop mRNA Decay. We used CRISPR/Cas9 to delete the Smr domain (Materials and Methods), generating *nonu-1(srf0780)*, which we hereafter refer to as *nonu-1(smrΔ)* (Figure 3D). Expression, splicing, and stability of the *nonu-1* transcript was not grossly perturbed in *nonu-1(smrΔ)* as assayed by RNA-seq. When combined with the *unc-54(Nonstop)* reporter, *nonu-1(smrΔ)* conferred derepression of GFP expression comparable to the *nonu-1* premature stop codon mutations originally isolated in the genetic screen (Figure 3E). We thus conclude that the NONU-1 Smr domain is required for Nonstop mRNA Decay.

We analyzed the sequence and structure of the NONU-1 Smr domain to better understand its potential catalytic mechanism. Sequence alignment of NONU-1 homologs across eukaryotes identified a highly conserved aspartate-x-histidine (DxH, where x is typically a hydrophobic amino acid) motif (Figure 3C, S3A) shared with two previously characterized endoribonucleases (Bhandari et al. 2011; Zhou et al. 2017). The DxH residues could conceivably function as catalytic residues. The DxH motif occupies a position in the Smr domain similar to the location of catalytic residues of other IF3-C-fold domains (Figure 3B). Given the defining DxH motif, we investigated if this motif is required for NONU-1’s role in Nonstop Decay. We generated alanine substitutions at this location (DxH>AxA). The resulting mutant (*nonu-1(AxA)*) exhibited a defect in Nonstop Decay comparable to the *nonu-1(smrΔ)* as well as *nonu-1* premature stop codon mutations (Figure 3E). Taken together, these observations are consistent with the idea that the DxH motif comprises the catalytic residues of the NONU-1 endoribonuclease.

It was recently reported that endonucleolytic cleavage during No-Go Decay occurs via a metal-independent nuclease resulting in a 3’phosphate and 5’hydroxl though the identity of this nuclease was not determined (Navickas et al.). The catalytic mechanism of Smr is unknown-- while purified Smr domain-containing proteins have cleavage activity on RNA, a catalytic mechanism has yet to be elucidated. However, the emerging picture of Smr endoribonuclease activity is consistent with what is known about cleavage during No-Go/Nonstop. First, *in vitro* RNA cleavage with Smr-domain containing proteins is inhibited by metals (Zhou et al. 2017). Second, metal-dependent nucleases typically require negatively charged amino acids (Asp or Glu) to chelate the positively charged metal. The one residue in NONU-1 that could conceivably function in this manner is Asp579 of the DxH motif. By itself this single Asp is unlikely to chelate a metal, though Asp579 could conceivably cooperate with other proteins and/or RNAs to create a metal-binding site.

To test whether Asp579 of the DxH motif could conceivably function as a metal chelator, we generated an isosteric mutant (D579N) by CRISPR/Cas9 that changes a single hydroxyl of the side chain to an amine. Aspartate to asparagine mutations in metal-dependent nucleases such as SMG-6 reduce metal binding and functionally eliminate RNA cleavage and SMG-6 function (Huntzinger et al. 2008). When combined with the *unc-54(Nonstop)* reporter, *nonu-1(D579N)* exhibited GFP expression similar to that of wild type *nonu-1* (Figure 3E). Functional substitution of Asp579 with an asparagine is most readily consistent with a metal-independent role for this residue in catalysis.

Our computational and experimental results are consistent with the model that NONU-1 is an endoribonuclease that acts during Nonstop mRNA Decay, with metal-independent cleavage being carried out by the highly conserved DxH motif within the Smr domain.

### NONU-1 reduces Nonstop mRNA levels

There are two main pathways known to repress protein expression from Nonstop mRNAs. One pathway is Ribosome Quality Control in which ubiquitination and CAT-tailing on the nascent peptidyl-tRNA cause protein aggregation and degradation (*e.g.*, (Bengtson and Joazeiro 2010; Shao et al. 2013; Shen et al. 2015)). There is also Nonstop mRNA Decay in which endo- and exo-nucleolytic digestion of the mRNA reduce mRNA levels and effectively prevent further rounds of translation (van Hoof et al. 2002; Schaeffer and van Hoof 2011). The *unc-54(Nonstop)* reporter contains a T2A ‘self-cleaving’ peptide after the open reading frame that allows the nascent peptide to leave the ribosome and escape repression from Ribosome Quality Control (Arribere and Fire 2018). That we identified *nonu-1* as a phenotypic suppressor of the *unc-54(Nonstop)* reporter suggests that *nonu-1* acts in Nonstop mRNA Decay rather than in Ribosome Quality Control (Figure 4A, S4). Consistent with this, by RNA-seq we detected a 2.4- fold increase of the *unc-54(Nonstop)* reporter mRNA in *nonu-1(smrΔ)* (Figure 4B, S5A). We thus conclude that *nonu-1* acts to reduce Nonstop mRNA levels.

**Figure 4:**
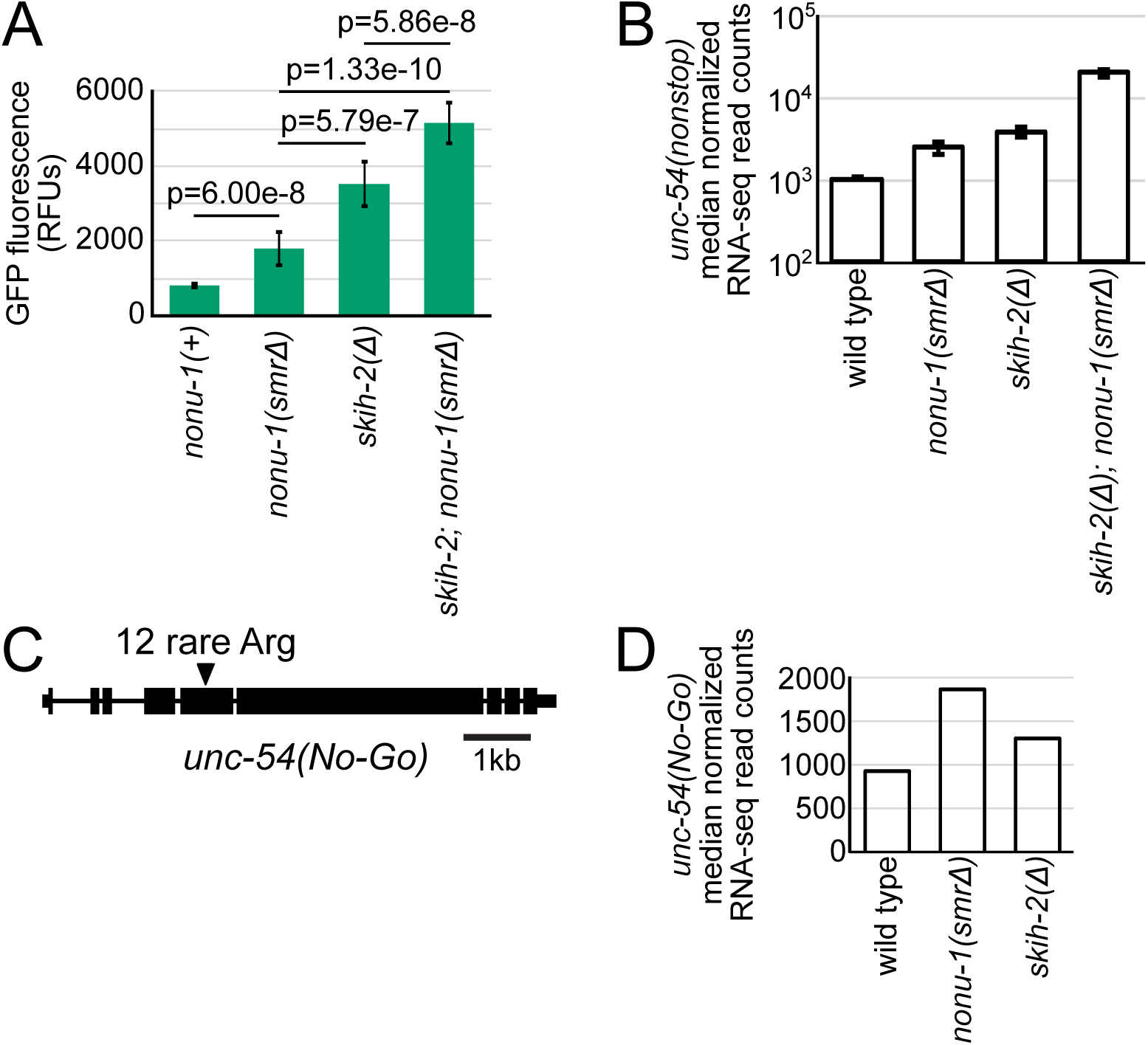
NONU-1 reduces mRNA levels of Nonstop and No-Go reporters. A. Quantification of GFP expression for the indicated strains (as in Figure 2C). *skih-2* encodes the catalytic subunit of an RNA helicase known to act in Nonstop mRNA Decay. P-values from Student’s T-test. RFUs stands for Relative Fluorescence Units. B. Median-normalized RNA-seq read counts (see Methods) for *unc-54(Nonstop)* across the indicated strains. Note axis is log-scaled. C. Gene diagram of *unc-54* No-Go Decay Reporter showing annotated exons (black rectangles) and introns (lines). Triangle indicates location of an insertion of 12 rare arginine codons (allele *srf0788*). Scale bar shows 1kb. D. Median-normalized RNA-seq read counts (see Methods) for *unc-54(No-Go)* across the indicated strains. Note axis is log-scaled.

We compared the phenotypic effect on *unc-54(Nonstop)* conferred by mutations in *nonu-1* relative to mutations in other Nonstop mRNA Decay components. Mutations in SKI complex subunits (*skih-2*, *ttc-37*) or ribosome rescue (*pelo-1*) suppress Nonstop mRNA Decay to different extents, likely reflecting their distinct functions in RNA stability and ribosome rescue (Arribere and Fire 2018; Guydosh and Green 2014; Shoemaker et al. 2010; Wilson et al. 2007). The phenotype of *nonu-1* mutants was distinct from *skih-2* null mutants, as assayed by the level of GFP fluorescence and *unc-54(Nonstop)* mRNA expression (Figure 4A, B). Thus NONU-1 is in a phenotypic class distinct from known Nonstop mRNA Decay suppressors.

Interestingly, the *skih-2 nonu-1* double mutant exhibited even greater Nonstop suppression than either single mutant alone, as assayed by Nonstop mRNA levels, Nonstop protein levels, and suppression of *unc-54*’s egg-laying phenotype (Figure 4A, B, S4). This result is consistent with the idea that phenotypic suppression by *skih-2* and *nonu-1* do not strictly depend on one another. It is unclear how much of the SKI complex’s repressive effect in Nonstop mRNA Decay is a direct result of accelerated mRNA decay versus other effects (*e.g.*, on mRNA translation initiation and/or recycling (Searfoss and Wickner 2000; Searfoss et al. 2001; Schmidt et al. 2016)).

Because the above analysis of *unc-54* mRNA expression was done by RNA-seq, we were able to address the question of whether *skih-2* and/or *nonu-1* are required for normal expression of endogenous mRNAs. There are endogenous mRNAs targeted by the Nonstop Decay pathway known in other organisms (*e.g.*, (Sparks and Dieckmann 1998)). Both *skih-2* and *nonu-1* are required for the normal expression of endogenous mRNAs: we observed that the mRNA levels from ∼20-40 genes increased in either *skih-2* or *nonu-1* (Methods, Supp Table S2,S3). While mRNAs that increased in a *skih-2* mutant also tended to increase in a *nonu-1* mutant (Figure S4B), our subsequent analyses support the idea that these mRNAs are not direct Nonstop targets. Many genes’ mRNAs that increased in either *skih-2* (13 of 27) or *nonu-1* (20 of 31) were derived from genes known to be developmentally regulated (Hendriks et al. 2014), consistent with a mild developmental delay between wild type and mutant strains. Furthermore, as a class, mRNAs whose levels increased in either *skih-2* or *nonu-1* did not exhibit any further increase in the double mutant (Figure S4B). This is in contrast to the behavior of the *unc-54(Nonstop)* reporter that showed increased mRNA levels in both single mutants (*skih-2* or *nonu-1*), and an additional increase in the double mutant (*skih-2 nonu-1*). We thus favor the idea that a majority of the endogenous mRNAs identified as upregulated in *skih-2* or *nonu-1* encode mRNAs that change as a result of secondary effects.

### NONU-1 acts in No-Go mRNA Decay

In addition to Nonstop Decay, endonucleolytic cleavage of the mRNA is thought to be an important step in No-Go Decay (Doma and Parker 2006). No-Go Decay results from blocks in translation elongation such as rare codons, polybasic amino acid stretches, and RNA structures (Doma and Parker 2006). We generated a No-Go Decay reporter in *C. elegans* by inserting 12 rare arginine codons in-frame in the *unc-54* gene (*unc-54(No-Go)*) (Figure 4C). We observed a two-fold derepression of the *unc-54(No-Go)* mRNA in *nonu-1(smrΔ)* mutant animals (Figure 4D). Thus NONU-1 is required for repression during No-Go Decay, and this function resides in the Smr domain. This result points to NONU-1 as a general player in translation surveillance.

We and others have recently shown that translation surveillance factors including *skih-2* and *pelo-1* act in NMD after a committed step of mRNA decay (Arribere and Fire 2018; Hashimoto et al. 2017). The effect of *skih-2* and *pelo-1* on NMD targets is not detectable at the level of RNA-seq, but is detectable as an accumulation of 15-18nt Ribo-seq footprints that represent a population of stalled ribosomes. Given its function in Nonstop Decay (Figures 2, 3), we expect that *nonu-1* also functions in NMD. As with *skih-2* and *pelo-1*, we failed to detect derepression of endogenous NMD targets by RNA-seq in *nonu-1* mutants. As it is not yet clear what population of ribosome (or di- or tri-ribosomes) is a substrate for NONU-1, it remains unclear how best to test for a role of *nonu-1* in NMD. Once there is additional information on the biochemistry, function, and relationship of NONU-1 to other translation surveillance events it will become possible to directly test the hypothesis that NONU-1 functions in NMD.

### Evolution of NONU-1 and Smr-domain proteins

Given that previous studies have only partly examined the evolution of Smr domains in eukaryotes (Liu et al. 2013), we conducted an in depth sequence and structure analysis of the Smr domain. Smr is an IF3-C fold domain, which also includes the nucleic acid-binding Alba, and the tRNA thiotransfer-catalyzing TusA (Figure 3B, C, S3A, S3B). One unifying sequence feature of this assemblage of IF3-C fold domains is a strongly-conserved sxs motif (where ‘s’ is a small residue, with the second ‘s’ typically a glycine) in the extended loop region between the second strand and the second helix thought to be involved in substrate-binding (Figure 3B, C, S3A, (Guo et al. 2014)). Among the catalytically active versions of the Smr-TusA-Alba assemblage, a conserved aspartate is observed at the C-terminal end of the initial core strand (Figure 3B, C, S3A). This aspartate is near a histidine in NONU-1 and forms the conserved DxH motif. More distant branches of the IF3-C fold include domains that bind, cleave, or process RNA, including the RNAseG/E nucleases (Fukui et al. 2008), the synaptojanin/calcineurin domain phosphoesterases and nucleases (Burroughs and Aravind 2016), the Schlafen domain endoribonucleases (Makarova et al. 2001; Li et al. 2018; Yang et al. 2018), and the RtcA RNA end cyclases (Figure S3B, (Palm et al. 2000)).

At least one NONU-1-like protein is traceable to the last eukaryotic common ancestor and is in practically all eukaryotic lineages, as would be expected of a core eukaryotic translation surveillance factor. NONU-1 homologs show a diversity of domain architectures across eukaryotes, including fusions to RNA-binding (CCCH, PWI), 2’-3’ cyclic phosphoesterase (2H), and ubiquitin binding and conjugation (UIM, UBL, and RING) domains (Figure S3C). We also noted multiple instances of rapidly-evolving lineage-specific expansions of NONU-1 homologs. These include multiple paralogs (25 or more) with distinct domain architectures in nematodes of the *Caenorhabditis* lineage. Some of these have predicted signal peptides suggesting that they are secreted (*e.g.*, in *Caenorhabditis remanei* and *Entamoeba*). The combination of lineage-specific expansions, rapid evolution, and secretion is a hallmark of proteins that are deployed as effectors in defensive or offensive roles in biological conflicts (Krishnan et al. 2018; Zhang et al. 2016; Lespinet et al. 2002). In light of this we hypothesize that several eukaryotic Smr proteins, especially the expanded versions, might function beyond translation surveillance as effectors deployed against viral or parasitic RNA. This is consistent with the discovery of a comparable role for the structurally related Schlafen domain in tRNA processing and retroviral RNA restriction (Li et al. 2012; Yang et al. 2018), as well as numerous studies showing that cellular RNA processing and translation surveillance factors have antiviral functions (Toh-E et al. 1978; Garcia et al. 2014; Balistreri et al. 2014).

## DISCUSSION

### The role of NONU-1 in translation surveillance

NONU-1 is the first factor with a nuclease domain to be identified as required for Nonstop/No-Go mRNA Decay. The Smr domain is conserved throughout eukaryotes and its function in translation surveillance is conserved between *S. cerevisiae* and *C. elegans*, establishing NONU-1 as an ancient factor critical to ribosome rescue and mRNA decay. Our identification and characterization of NONU-1 sheds light on the poorly understood intersection of translation and mRNA decay and sets the stage for a more complete molecular understanding of ribosome rescue.

Interestingly, we note that NONU-1 is not required for full repression of the mRNA targets of translation surveillance. Even in presumed *nonu-1* null mutants we observed substantial repression of Nonstop/No-Go targets that could be relieved with other suppressors (*e.g.*, *skih-2*). Two simple, non-mutually exclusive models are: (1) NONU-1 may function redundantly with other endonucleases in translation surveillance. Recent work in *S. cerevisiae* points to the existence of at least two nucleases active during No-Go Decay though their identities remain unknown (Ikeuchi et al. 2019). (2) There may be cleavage-independent mechanisms that repress the mRNA targets of translation surveillance. Whether through additional nucleases or cleavage-independent mechanisms, our work supports a redundancy in translation surveillance that ensures robust repression of its targets and efficient rescue of stalled ribosomes.

Our results support the idea that NONU-1 acts in translation surveillance largely independently of the SKI complex. We observed multiplicative effects in *skih-2 nonu-1* double mutants, and in both *C. elegans* and *S. cerevisiae* we observed a greater derepression of Nonstop reporters in *skih-2*/*ski2Δ* mutants relative to *nonu-1*/*cue2Δ*/*ypl199cΔ* mutants. This is surprising given the prevailing model in the field that SKI accelerates 3’>5’ decay after endonucleolytic cleavage. One possibility is that NONU-1 may function redundantly with other endonucleases to create SKI substrates. Another possibility is that SKI’s role in surveillance is misunderstood. While it is widely known that one function of the SKI complex is to destabilize the upstream (5’) mRNA fragment during translation surveillance (*e.g.*, (Doma and Parker 2006; Hashimoto et al. 2017)), it is unclear if this effect is responsible for the phenotypic suppression of Nonstop reporters by SKI. Alternative models include functional suppression by SKI’s effects on ribosome recycling or initiation (Searfoss and Wickner 2000; Searfoss et al. 2001), which is also consistent with recent structural data showing a direct role for SKI on the ribosome and near the 5’ends of ORFs (Schmidt et al. 2016).

In *S. cerevisiae* endonucleolytic cleavage during No-Go creates a 5’hydroxyl that is phosphorylated by Rlg1/Trl1 to facilitate 5’>3’ digestion by Xrn1 (Navickas et al.). While Rlg1/Trl1 is widely conserved in several eukaryotic lineages including fungi, plants, alveolates, and kinetoplastids (Burroughs and Aravind 2016), its absence in the animal/choanoflagellate lineage raises the question of what protein carries out this RNA repair. Interestingly, the animal/choanoflagellate NONU-1 homologs have acquired a P-loop kinase domain that is related to but distinct from the kinase domain of Rlg1/Trl1. The acquisition of a P-loop kinase domain in NONU-1 in organisms that have lost Rlg1/Trl1 suggests a model in which NONU-1 phosphorylates its own cleavage products to facilitate degradation by Xrn1.

## ACKNOWLEDGEMENTS

We thank Manny Ares, Grant Hartzog, and their respective labs for providing yeast strains, reagents, and expertise; Ben Abrams and the UCSC Microscopy Facility for imaging and quantification; Beth Shapiro, Beth Nelson, the UCSC Paleogenomics Lab, and the Vincent J. Coates Genomics Sequencing Facility for deep sequencing; Ambro van Hoof for sharing pAV188 (*his3::nonstop* reporter plasmid); Susan Strome and lab for reagents and microscopy early on in the project. We thank Manny Ares, Susan Strome, Al Zahler, and Sara Dubbury for comments on the manuscript. This work was supported in part by the Intramural Research Program of the NIH, National Library of Medicine (AMB and LA), an R01 from NIGMS at the NIH awarded to JAA (1R01GM131012-01), and start-up funds from UCSC awarded to JAA.

## AUTHOR CONTRIBUTIONS

MLG and JAA conceived of the study and designed, conducted, and analyzed experiments. TAE performed RNA-seq and genomic sequencing. MNP isolated several *nonu-1* alleles. AMB and LA performed *in silico* protein analyses, some of which informed additional experiments carried out by MLG and JAA. The manuscript was written by MLG and JAA with some sections by AMB and LA and contributions from all authors.

## DATA AVAILABILITY

SRA number is pending.

## MATERIALS & METHODS

### *C. elegans* growth and propagation

Strains were derived from N2 background (VC2010, (Thompson et al. 2013)). Animals were grown at 22C on NGM plates using OP50 as a food source per standard *C. elegans* husbandry (Brenner 1974). Some strains were provided by the CGC, which is funded by NIH Office of Research Infrastructure Programs (P40 OD010440).

### EMS Mutagenesis

EMS mutagenesis was performed essentially as described (Arribere and Fire 2018). Briefly, a large population of *unc-54(cc4092)* was washed with M9 and resuspended in a final volume of 4ml M9. EMS was added to a final concentration of ∼1mM and animals incubated for 4 hours at room temperature with rotation. Animals were washed and allowed to recover overnight on plates with OP50. The next day animals were washed and eggs isolated via sodium hypochlorite treatment. 100-200 eggs were placed on a single small NGM plate and allowed to develop. Plates were screened for individuals with increased GFP fluorescence at the F2/F3 generation. Only a single isolate was kept per small NGM plate, ensuring independence of mutations identified.

### Mutation mapping

We crossed each isolated suppressor to Hawaiian *unc-54(cc4112)* males (expressing an UNC-54::mCherry fusion engineered by CRISPR/Cas9). Cross progeny were picked to new plates to self fertilize. From among the F2, we picked ∼30 GFP+ progeny to a new plate and allowed the animals to self-fertilize and starve. Upon starvation, animals were washed off the plate with 1mL EN50, and further washed with EN50 to remove residual *E. coli*. Genomic DNA was extracted after proteinase K treatment and resuspended in 50uL TE pH7.4. 50ng genomic DNA was used as input for Nextera (Tn5) sequencing libraries. Libraries were sequenced at the Vincent J. Coates Genomics Sequencing Laboratory at UC Berkeley.

Reads were mapped to the *C. elegans* genome using bowtie2 (version 2.3.4.1). Reads were assigned using GATK (McKenna et al. 2010) and a previously published dataset of Hawaiian SNPs (Thompson et al. 2013). The fraction of reads that were assignable to Hawaiian or N2 animals was calculated across the genome, and linkage was identified by portions of the genome that went to 0% Hw. Regions of linkage were then manually inspected to identify candidate lesions/loci.

### CRISPR/Cas9 mutagenesis

CRISPR/Cas9 was performed as previously described (Arribere et al. 2014). A full list of gRNAs is available in Table S1, and exact sequences of mutant alleles is provided alongside *C. elegans* strains in Table S1. Multiple genetic isolates of each mutation were obtained and observed to have identical phenotypes.

### Microscopy and Image Quantification

L4 worms were anesthetized in 2uL 1mM levamisole in a microscope well slide with a .15mm coverslip. A Zeiss AxioZoom microscope was used with a 1.0x objective and a GFP fluorescence light source to acquire all images. The *unc-54(cc4092); skih-2(cc2854)* strain was used to set parameters (exposure time 330ms., shift 50%, zoom 80%) and the same parameters were used for all images. Five representative worms were imaged for each strain. All comparisons shown are between images obtained during the same imaging session.

We used FIJI to define area of the worm, subtract the background, and determine mean pixel intensity for the area of each worm. A mean fluorescence intensity was calculated for each strain. We calculated two standard deviations above and below the mean to obtain a 95% confidence intervals.

### RNA-seq and Analysis

25-50 day 1 adults were picked from a blank plate into S-basal solution and washed to remove *E. coli*. Animals were dissolved in trizol and total RNA extracted. Ribosomal RNA was depleted from 250 ng of total RNA using an NEBNext rRNA Depletion Kit (Human/Mouse/Rat) (catalog number E6310) and libraries were constructed using an NEBNext Ultra RNA Library Prep Kit for Illumina sequencing (catalog number E7530). Libraries were sequenced at the University of California, Santa Cruz using the Illumina NextSeq platform.

RNA-seq reads were trimmed with cutadapt and mapped using STAR (version 2.5.0a) to the *C. elegans* genome (WBCel235) with the *unc-54* locus modified to match the *unc-54(cc4092)* allele. Reads that mapped within the annotated bounds of a protein coding gene were assigned to that gene. Multiply-mapping reads or reads that could not be unambiguously assigned to a gene (*e.g.*, due to overlapping genes) were discarded. Read counts were median-normalized using DESeq (Anders and Huber 2010).

For differential expression of endogenous mRNAs in *skih-2* and *nonu-1*, genes with mRNAs that increased in biological duplicates (*skih-2*) or triplicates (*nonu-1*) were identified with DESeq. mRNA levels were deemed significantly different if they exhibited an adjusted p-value <0.05 (*skih-2*) or <0.001 (*nonu-1*). Varying these cutoffs changed the number genes identified as *skih-2* or *nonu-1* targets, but did not alter our conclusions.

### S. cerevisiae husbandry

*S. cerevisiae* strains were grown on YPAD media at 30C. A full list of strains, sources, and constructions is available in Table S1. All mutations were verified by at least two independent PCR primers.

Cells were transformed with the *his3::nonstop* reporter plasmid (pAV188) via lithium acetate transformation and plated on selective media (SD-Ura). Two Ura+ transformants were taken for each strain, and results were reproducible across these independent isolates. Cells were subsequently grown on SD-Ura plates and in SD-Ura liquid media to maintain the plasmid. For the *his3::nonstop* reporter assay, 5ml liquid cultures were grown overnight to saturation. The OD600 was measured and the same OD600 was used as the starting number of cells for all strains. Cells were serially diluted 1:6 and plated on selective media (SD-Ura and SD-Ura-His). Plates were photographed after ∼8 days of growth at 30C.

### Sequence Analysis

Domain sequence similarity searches were performed using PSI-BLAST program (Altschul et al. 1997) against the non-redundant (nr) database housed at the NCBI and the HHpred program (Zimmermann et al. 2018) against pfam and pdb databases (Finn et al. 2016; Burley et al. 2019). Structure similarity searches were performed using the DALI server (Holm and Sander 1995). Multiple sequence alignments were built using the Kalign2 program, with manual adjustments based on profile-profile and high-scoring pair sequence similarity search results (Lassmann et al. 2009). Domain architectures Smr domain-containing proteins were elucidated by first running rpsblast searches against a PSSM library constructed from the pfam profile database (Marchler-Bauer and Bryant 2004). Regions lacking any annotation were then used as seeds in further rounds of iterative similarity searches. Phylogenetic analyses were carried out using approximate-maximum-likelihood as enacted by the FastTree 2.1 program with default parameters for amino acid sequences (Price et al. 2010).

## SUPPLEMENTAL FIGURES

**Figure S1:**
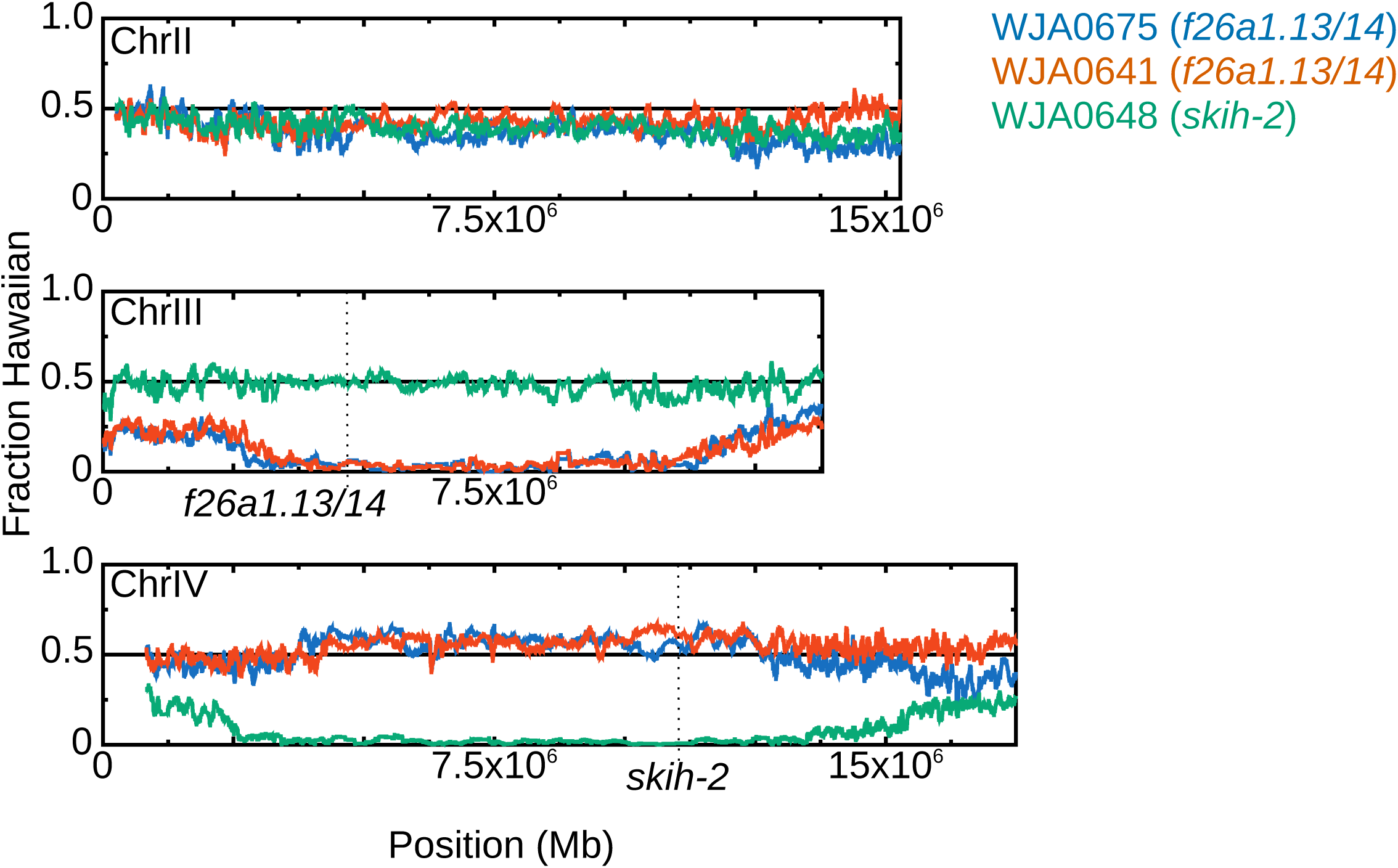
Two strains show linkage to an area of chromosome III lacking known Nonstop mRNA Decay factors. Genetic maps for three strains showing linkage (and lack thereof) to chromosomes II, III, and IV. WJA0648 contains a mutation in *skih-2* on chromosome IV. WJA0675 and WJA0641 contain mutations in *f26a1.13/14* (renamed *nonu-1*) on chromosome III. X-axis shows position along the chromosome in megabases (Mb). See methods for a description of the Hawaiian mapping procedure and variant calling. This technique narrows down the causative mutation to a large swath of one chromosome, after which we manually inspected the region of interest to identify possible mutations (subsequently verified by CRISPR/Cas9).

**Figure S2:**
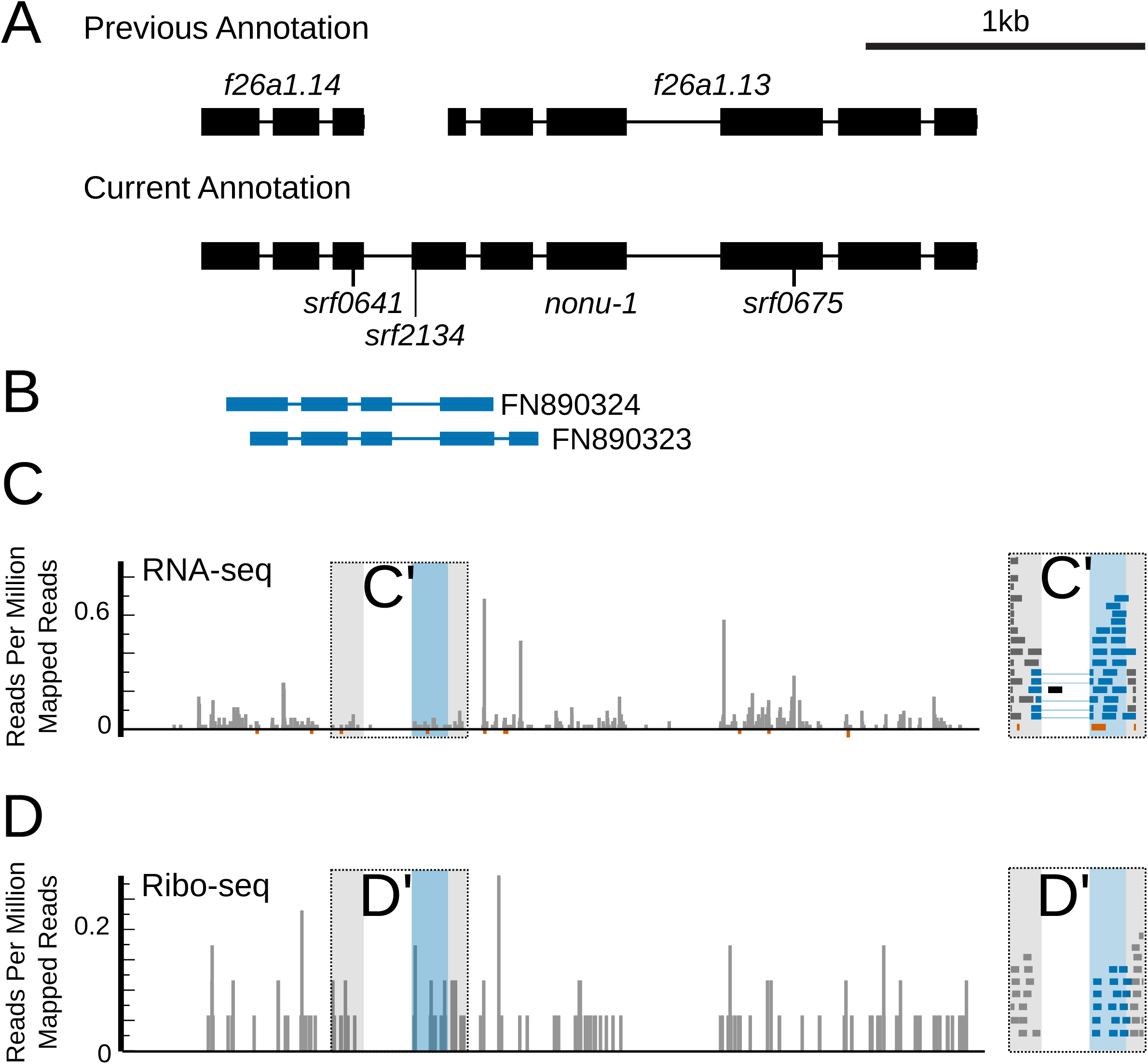
*f26a1.13/14* encode a single functional gene. A. Gene diagrams for *f26a1.13* and *f26a1.14*. Prior annotations indicated on top, with rectangles and lines representing exons and introns, respectively. New annotation below, along with the location of each of three premature stop codon mutations (*srf* alleles) that exhibit identical phenotypes in a Nonstop mRNA reporter strain. Data in parts (B-D) are vertically aligned with annotations in part (A). B. Public ESTs that span the proposed novel splice junction. C. Published RNA-seq data from (Hendriks et al., 2014) showing reads spanning the proposed novel splice junction. Inset (C’) zooms in on the region of interest, with reads supporting the junction highlighted (blue). Antisense reads in red. D. Published Ribo-seq data from (Hendriks et al., 2014) showing ribosome footprints spanning the proposed novel splice junction. Inset (D’) zooms in on the region of interest, with reads supporting the junction highlighted (blue). Note there is a similar density of Ribo-seq reads throughout the entire gene body of both *f26a1.13* and *f26a1.14*, consistent with the idea they represent a single translational unit.

**Figure S3:**
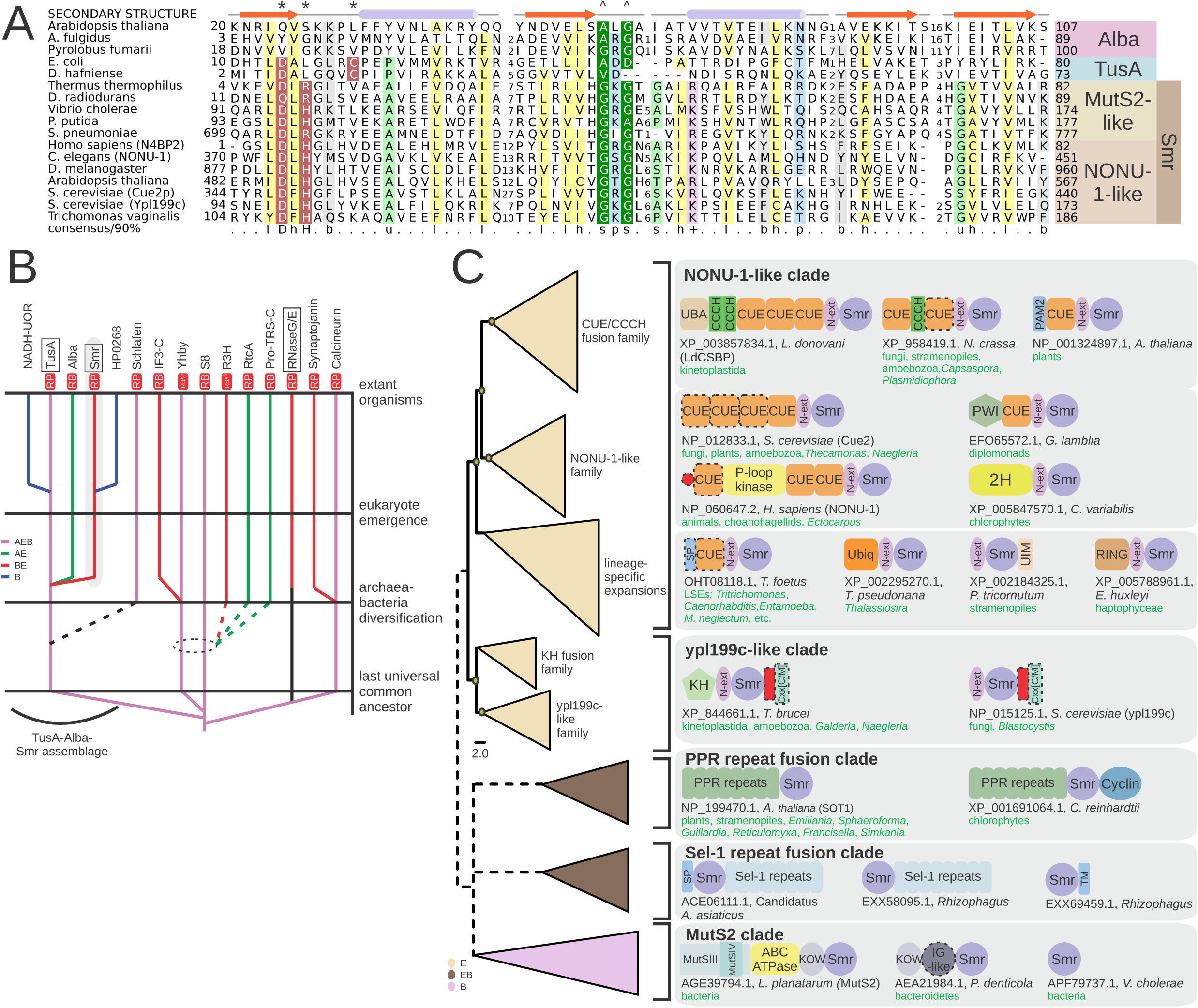
Conservation and evolution of the Smr domain in the IF3-C fold framework. A. Extended multiple sequence alignment of representatives of domains from the TusA-Alba-Smr assemblage. Secondary structure shown on top, and consensus on which alignment is colored provided on bottom line. B. Phylogenetic tree of evolutionary relationships between different IF3-C fold domains, with key evolutionary transitions labeled to the left of the diagram. Tree branches are colored according to occurrence of the domain in the respective kingdom (Archaea, Bacteria, and/or Eukarya). Some evolutionary relationships are uncertain (dotted lines). Structural renderings of boxed domains (TusA, Smr, RNaseG/E) are shown in Figure 2D. Functional annotations for lineages in red boxes below names are as follows: ‘RB’, RNA-binding; ‘RP’, RNA-processing. Here, top-down view of RNaseG/E is provided to emphasize the importance of dimerization in the metal-binding nuclease active site. C. Broader phylogeny depicted as stylized tree of all known Smr domain families. Nodes supported by bootstrap >75% are marked by brown circles. Dotted lines in tree indicate uncertainty in phylogenetic placement. Representative domain architectures observed within each family provided to the right of monophyletic, collapsed tree branches. Individual domains found within a single polypeptide are represented by distinct shapes labeled by name. Dotted lines around a domain indicate absence in some organisms of the listed phylogenetic distribution for each architecture, provided below architectural depiction. Red circle represents the N-terminal basic stretch described in the text. Red box represents a variable-length charged region characteristic of members of the ypl199c-like clade.

**Figure S4:**
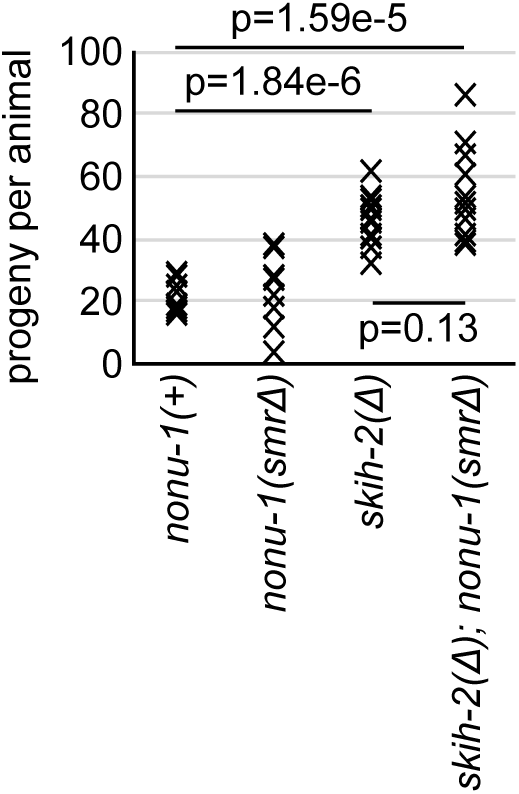
*skih-2* and *skih-2;nonu-1* have increased brood sizes. Brood sizes for the indicated strains. Each X represents the number of progeny from a single animal from that strain. 12 animals were examined per strain. P-values from Student’s T-test.

**Figure S5:**
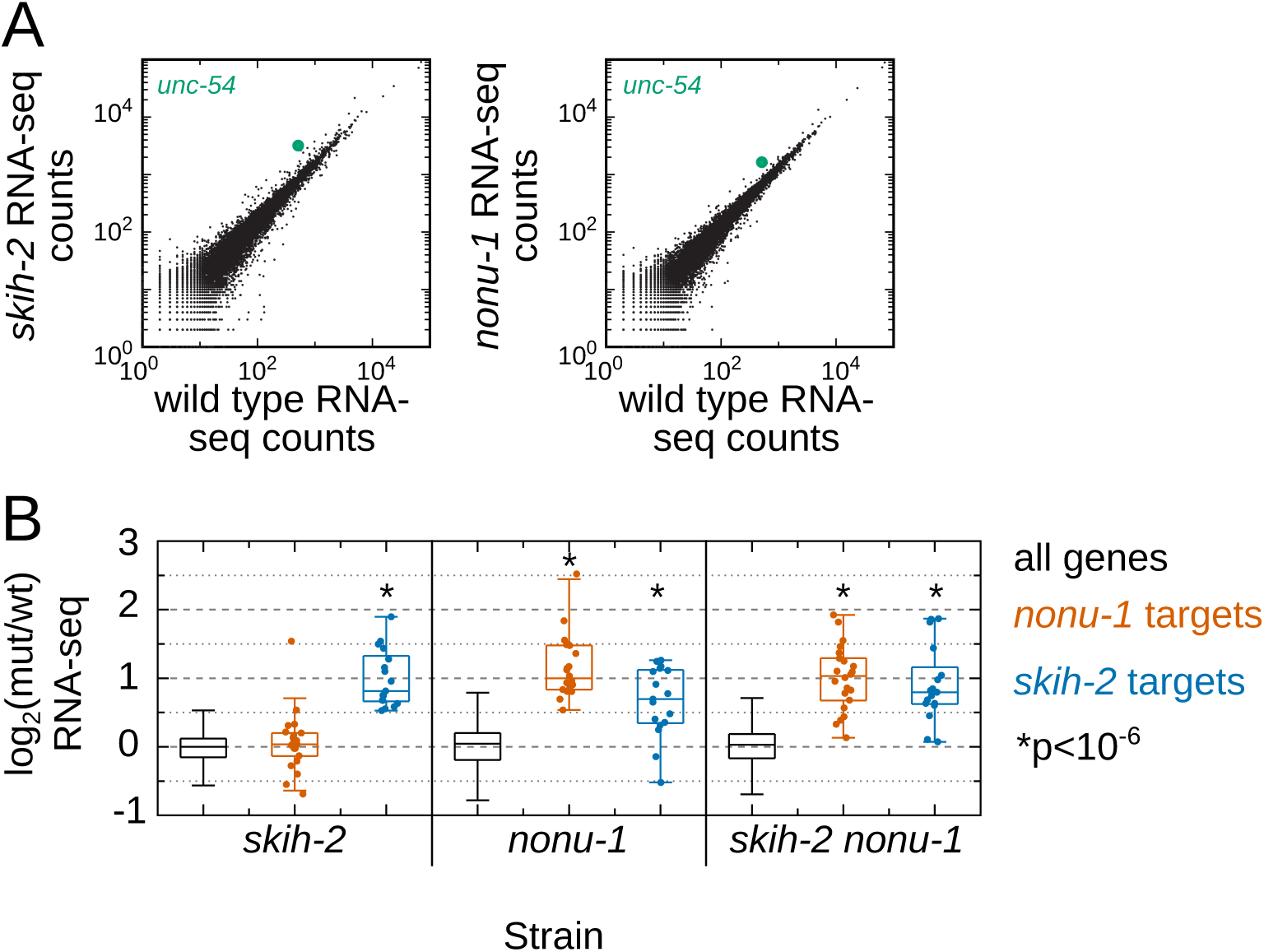
Endogenous mRNAs regulated by *skih-2* or *nonu-1* are likely indirect targets. A: RNA-seq read counts in the indicated strains. RNA-seq was performed (see Methods), and read counts for all mRNAs is shown (black dots) with *unc-54* highlighted (green dot). Off-diagonal genes indicate increased mRNA expression in one strain relative to the other. Note axes are log-scaled. B: Y-axis shows log2 fold change for gene expression changes between the indicated mutant strains and wild type for all genes with at least 30 reads in wild type (different read cutoffs in wild type yielded similar results). *nonu-1* (red) and *skih-2* (blue) targets are defined as mRNAs that are upregulated in the respective mutant strains relative to wild type (as determined by DESeq, see Methods). P-value from Mann Whitney U test comparing the indicated subset of genes to all genes. Note that unlike *unc-54*, targets of *nonu-1* and *skih-2* fail to exhibit a further increase in mRNA expression in the double mutant. A biological replicate of these data produced similar results.

## SUPPLEMENTARY TABLES

**Supplementary Table 1 (attached):** *C. elegans* strains, *S. cerevisiae* strains, and oligonucleotides used in this study.

**Supplementary Table 2:**
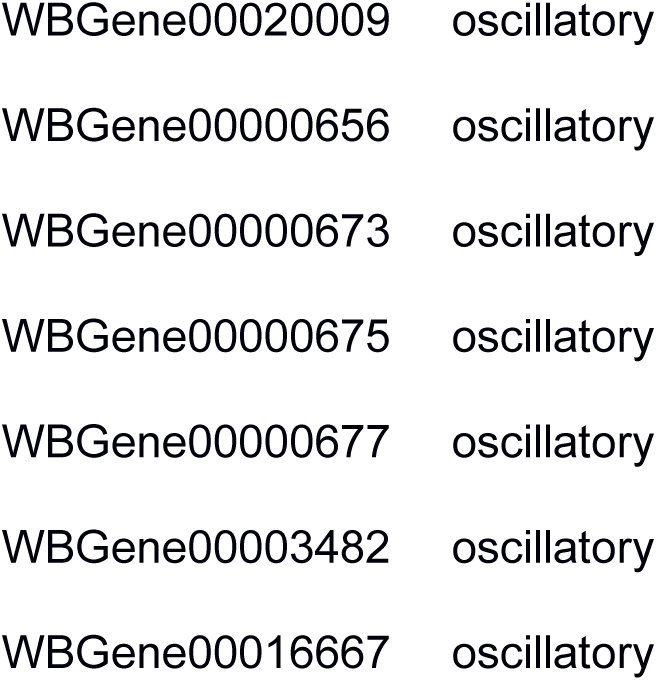
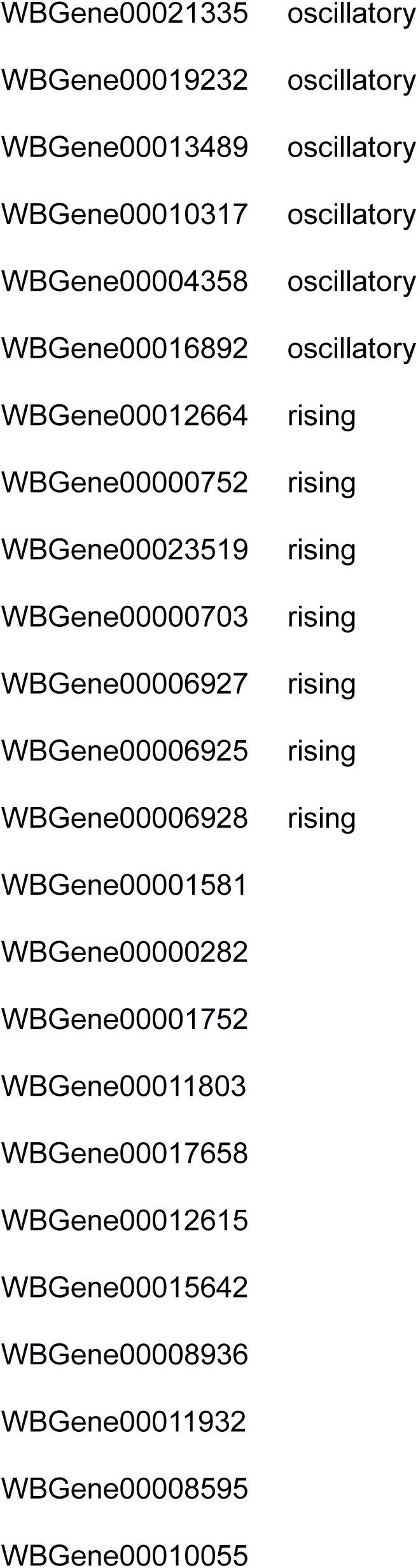
Genes with mRNAs significantly increased in *nonu-1* (p_adj<0.001) and their annotated developmental regulation in (Hendriks et al., 2014)

**Supplementary Table 3:**
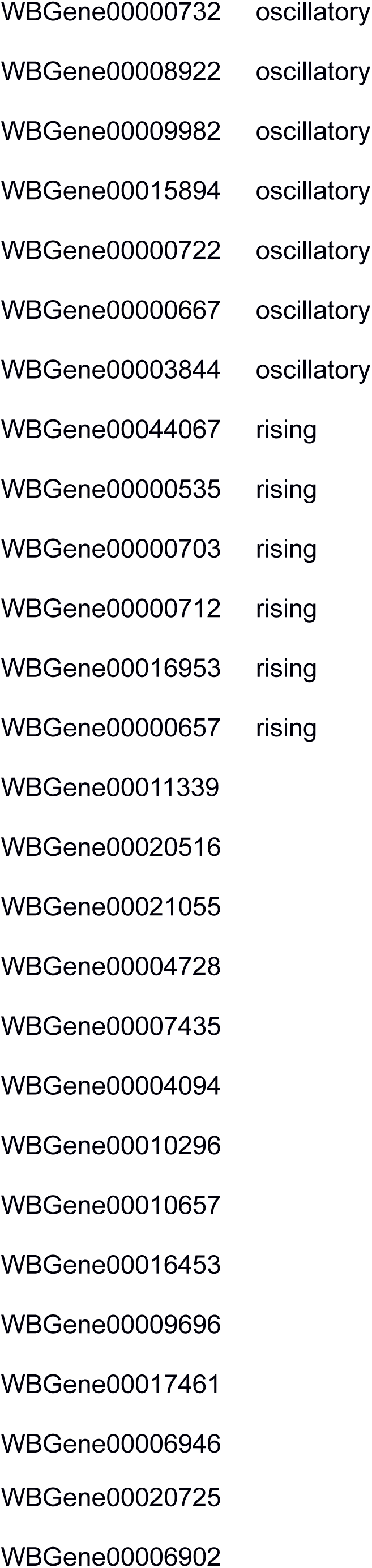
Genes with mRNAs significantly increased in *skih-2* (p_adj<0.05) and their annotated developmental regulation in (Hendriks et al., 2014)

## REFERENCES

Altschul SF, Madden TL, Schäffer AA, Zhang J, Zhang Z, Miller W, Lipman DJ. 1997. Gapped BLAST and PSI-BLAST: a new generation of protein database search programs. Nucleic Acids Res 25: 3389–3402.

Anderson P, Brenner S. 1984. A selection for myosin heavy chain mutants in the nematode Caenorhabditis elegans. Proc Natl Acad Sci U S A 81: 4470–4474.

Anders S, Huber W. 2010. Differential expression analysis for sequence count data. Genome Biol 11: R106.

Aravind L, Iyer LM, Anantharaman V. 2003. The two faces of Alba: the evolutionary connection between proteins participating in chromatin structure and RNA metabolism. Genome Biol 4: R64.

Arribere JA, Bell RT, Fu BXH, Artiles KL, Hartman PS, Fire AZ. 2014. Efficient marker-free recovery of custom genetic modifications with CRISPR/Cas9 in Caenorhabditis elegans. Genetics 198: 837–846.

Arribere JA, Fire AZ. 2018. Nonsense mRNA suppression via Nonstop decay. Elife 7. http://dx.doi.org/10.7554/eLife.33292.

Balistreri G, Horvath P, Schweingruber C, Zünd D, McInerney G, Merits A, Mühlemann O, Azzalin C, Helenius A. 2014. The host nonsense-mediated mRNA decay pathway restricts Mammalian RNA virus replication. Cell Host Microbe 16: 403–411.

Bejsovec A, Anderson P. 1988. Myosin heavy-chain mutations that disrupt Caenorhabditis elegans thick filament assembly. Genes Dev 2: 1307–1317.

Bengtson MH, Joazeiro CAP. 2010. Role of a ribosome-associated E3 ubiquitin ligase in protein quality control. Nature 467: 470–473.

Bhandari D, Guha K, Bhaduri N, Saha P. 2011. Ubiquitination of mRNA cycling sequence binding protein from Leishmania donovani (LdCSBP) modulates the RNA endonuclease activity of its Smr domain. FEBS Lett 585: 809–813.

Brenner S. 1974. The genetics of Caenorhabditis elegans. Genetics 77: 71–94.

Burley SK, Berman HM, Bhikadiya C, Bi C, Chen L, Di Costanzo L, Christie C, Dalenberg K, Duarte JM, Dutta S, et al. 2019. RCSB Protein Data Bank: biological macromolecular structures enabling research and education in fundamental biology, biomedicine, biotechnology and energy. Nucleic Acids Res 47: D464–D474.

Burroughs AM, Aravind L. 2016. RNA damage in biological conflicts and the diversity of responding RNA repair systems. Nucleic Acids Res 44: 8525–8555.

Dibb NJ, Brown DM, Karn J, Moerman DG, Bolten SL, Waterston RH. 1985. Sequence analysis of mutations that affect the synthesis, assembly and enzymatic activity of the unc-54 myosin heavy chain of Caenorhabditis elegans. J Mol Biol 183: 543–551.

Dibb NJ, Maruyama IN, Krause M, Karn J. 1989. Sequence analysis of the complete Caenorhabditis elegans myosin heavy chain gene family. J Mol Biol 205: 603–613.

Doitsidou M, Poole RJ, Sarin S, Bigelow H, Hobert O. 2010. C. elegans mutant identification with a one-step whole-genome-sequencing and SNP mapping strategy. PLoS One 5: e15435.

Doma MK, Parker R. 2006. Endonucleolytic cleavage of eukaryotic mRNAs with stalls in translation elongation. Nature 440: 561–564.

Epstein HF, Waterston RH, Brenner S. 1974. A mutant affecting the heavy chain of myosin in Caenorhabditis elegans. J Mol Biol 90: 291–300.

Finn RD, Coggill P, Eberhardt RY, Eddy SR, Mistry J, Mitchell AL, Potter SC, Punta M, Qureshi M, Sangrador-Vegas A, et al. 2016. The Pfam protein families database: towards a more sustainable future. Nucleic Acids Res 44: D279–85.

Fukui K, Nakagawa N, Kitamura Y, Nishida Y, Masui R, Kuramitsu S. 2008. Crystal structure of MutS2 endonuclease domain and the mechanism of homologous recombination suppression. J Biol Chem 283: 33417–33427.

Garcia D, Garcia S, Voinnet O. 2014. Nonsense-mediated decay serves as a general viral restriction mechanism in plants. Cell Host Microbe 16: 391–402.

Garzia A, Jafarnejad SM, Meyer C, Chapat C, Gogakos T, Morozov P, Amiri M, Shapiro M, Molina H, Tuschl T, et al. 2017. The E3 ubiquitin ligase and RNA-binding protein ZNF598 orchestrates ribosome quality control of premature polyadenylated mRNAs. Nat Commun 8: 16056.

Guo L, Ding J, Guo R, Hou Y, Wang D-C, Huang L. 2014. Biochemical and structural insights into RNA binding by Ssh10b, a member of the highly conserved Sac10b protein family in Archaea. J Biol Chem 289: 1478–1490.

Guydosh NR, Green R. 2014. Dom34 rescues ribosomes in 3’ untranslated regions. Cell 156: 950–962.

Hashimoto Y, Takahashi M, Sakota E, Nakamura Y. 2017. Nonstop-mRNA decay machinery is involved in the clearance of mRNA 5’-fragments produced by RNAi and NMD in Drosophila melanogaster cells. Biochem Biophys Res Commun 484: 1–7.

Hendriks G-J, Gaidatzis D, Aeschimann F, Großhans H. 2014. Extensive oscillatory gene expression during C. elegans larval development. Mol Cell 53: 380–392.

Hodgkin J, Papp A, Pulak R, Ambros V, Anderson P. 1989. A new kind of informational suppression in the nematode Caenorhabditis elegans. Genetics 123: 301–313.

Holm L, Sander C. 1995. Dali: a network tool for protein structure comparison. Trends Biochem Sci 20: 478–480.

Huntzinger, Eric, Isao Kashima, Maria Fauser, Jérôme Saulière, and Elisa Izaurralde. 2008. “SMG6 Is the Catalytic Endonuclease That Cleaves mRNAs Containing Nonsense Codons in Metazoan.” RNA 14 (12): 2609–17.

Ikeuchi K, Tesina P, Matsuo Y, Sugiyama T, Cheng J, Saeki Y, Tanaka K, Becker T, Beckmann R, Inada T. 2019. Collided ribosomes form a unique structural interface to induce Hel2-driven quality control pathways. EMBO J 38. http://dx.doi.org/10.15252/embj.2018100276.

Joazeiro CAP. 2017. Ribosomal Stalling During Translation: Providing Substrates for Ribosome-Associated Protein Quality Control. Annu Rev Cell Dev Biol 33: 343–368.

Juszkiewicz S, Chandrasekaran V, Lin Z, Kraatz S, Ramakrishnan V, Hegde RS. 2018. ZNF598 Is a Quality Control Sensor of Collided Ribosomes. Mol Cell 72: 469–481.e7.

Kang RS, Daniels CM, Francis SA, Shih SC, Salerno WJ, Hicke L, Radhakrishnan I. 2003. Solution structure of a CUE-ubiquitin complex reveals a conserved mode of ubiquitin binding. Cell 113: 621–630.

Krishnan A, Iyer LM, Holland SJ, Boehm T, Aravind L. 2018. Diversification of AID/APOBEC-like deaminases in metazoa: multiplicity of clades and widespread roles in immunity. Proc Natl Acad Sci U S A 115: E3201–E3210.

Lassmann T, Frings O, Sonnhammer ELL. 2009. Kalign2: high-performance multiple alignment of protein and nucleotide sequences allowing external features. Nucleic Acids Res 37: 858–865.

Lee RYN, Howe KL, Harris TW, Arnaboldi V, Cain S, Chan J, Chen WJ, Davis P, Gao S, Grove C, et al. 2018. WormBase 2017: molting into a new stage. Nucleic Acids Res 46: D869–D874.

Leipe DD, Koonin EV, Aravind L. 2003. Evolution and classification of P-loop kinases and related proteins. J Mol Biol 333: 781–815.

Lespinet O, Wolf YI, Koonin EV, Aravind L. 2002. The role of lineage-specific gene family expansion in the evolution of eukaryotes. Genome Res 12: 1048–1059.

Li M, Kao E, Gao X, Sandig H, Limmer K, Pavon-Eternod M, Jones TE, Landry S, Pan T, Weitzman MD, et al. 2012. Codon-usage-based inhibition of HIV protein synthesis by human schlafen 11. Nature 491: 125–128.

Li M, Kao E, Malone D, Gao X, Wang JYJ, David M. 2018. DNA damage-induced cell death relies on SLFN11-dependent cleavage of distinct type II tRNAs. Nat Struct Mol Biol 25: 1047–1058.

Liu S, Melonek J, Boykin LM, Small I, Howell KA. 2013. PPR-SMRs: ancient proteins with enigmatic functions. RNA Biol 10: 1501–1510.

Makarova KS, Aravind L, Wolf YI, Tatusov RL, Minton KW, Koonin EV, Daly MJ. 2001. Genome of the extremely radiation-resistant bacterium Deinococcus radiodurans viewed from the perspective of comparative genomics. Microbiol Mol Biol Rev 65: 44–79.

Marchler-Bauer A, Bryant SH. 2004. CD-Search: protein domain annotations on the fly. Nucleic Acids Res 32: W327–31.

McKenna A, Hanna M, Banks E, Sivachenko A, Cibulskis K, Kernytsky A, Garimella K, Altshuler D, Gabriel S, Daly M, et al. 2010. The Genome Analysis Toolkit: a MapReduce framework for analyzing next-generation DNA sequencing data. Genome Res 20: 1297–1303.

Moerman DG, Plurad S, Waterston RH, Baillie DL. 1982. Mutations in the unc-54 myosin heavy chain gene of Caenorhabditis elegans that alter contractility but not muscle structure. Cell 29: 773–781.

Navickas A, Chamois S, Saint-Fort R, Henri J, Torchet C, Benard L. A unique No-Go Decay cleavage in mRNA exit-tunnel of ribosome produces 5′-OH ends phosphorylated by Rlg1. http://dx.doi.org/10.1101/465633.

Palm GJ, Billy E, Filipowicz W, Wlodawer A. 2000. Crystal structure of RNA 3’-terminal phosphate cyclase, a ubiquitous enzyme with unusual topology. Structure 8: 13–23.

Price MN, Dehal PS, Arkin AP. 2010. FastTree 2--approximately maximum-likelihood trees for large alignments. PLoS One 5: e9490.

Saito K, Horikawa W, Ito K. 2015. Inhibiting K63 polyubiquitination abolishes no-go type stalled translation surveillance in Saccharomyces cerevisiae. PLoS Genet 11: e1005197.

Schaeffer D, van Hoof A. 2011. Different nuclease requirements for exosome-mediated degradation of normal and nonstop mRNAs. Proc Natl Acad Sci U S A 108: 2366–2371.

Schmidt C, Kowalinski E, Shanmuganathan V, Defenouillère Q, Braunger K, Heuer A, Pech M, Namane A, Berninghausen O, Fromont-Racine M, et al. 2016. The cryo-EM structure of a ribosome-Ski2-Ski3-Ski8 helicase complex. Science 354: 1431–1433.

Searfoss A, Dever TE, Wickner R. 2001. Linking the 3’ poly(A) tail to the subunit joining step of translation initiation: relations of Pab1p, eukaryotic translation initiation factor 5b (Fun12p), and Ski2p-Slh1p. Mol Cell Biol 21: 4900–4908.

Searfoss AM, Wickner RB. 2000. 3’ poly(A) is dispensable for translation. Proc Natl Acad Sci U S A 97: 9133–9137.

Shao S, von der Malsburg K, Hegde RS. 2013. Listerin-dependent nascent protein ubiquitination relies on ribosome subunit dissociation. Mol Cell 50: 637–648.

Shen PS, Park J, Qin Y, Li X, Parsawar K, Larson MH, Cox J, Cheng Y, Lambowitz AM, Weissman JS, et al. 2015. Protein synthesis. Rqc2p and 60S ribosomal subunits mediate mRNA-independent elongation of nascent chains. Science 347: 75–78.

Shoemaker CJ, Eyler DE, Green R. 2010. Dom34:Hbs1 promotes subunit dissociation and peptidyl-tRNA drop-off to initiate no-go decay. Science 330: 369–372.

Simms CL, Yan LL, Zaher HS. 2017. Ribosome Collision Is Critical for Quality Control during No-Go Decay. Mol Cell 68: 361–373.e5.

Sparks KA, Dieckmann CL. 1998. Regulation of poly(A) site choice of several yeast mRNAs. Nucleic Acids Res 26: 4676–4687.

Thompson O, Edgley M, Strasbourger P, Flibotte S, Ewing B, Adair R, Au V, Chaudhry I, Fernando L, Hutter H, et al. 2013. The million mutation project: a new approach to genetics in Caenorhabditis elegans. Genome Res 23: 1749–1762.

Toh-E A, Guerry P, Wickner RB. 1978. Chromosomal superkiller mutants of Saccharomyces cerevisiae. J Bacteriol 136: 1002–1007.

van Hoof A, Frischmeyer PA, Dietz HC, Parker R. 2002. Exosome-mediated recognition and degradation of mRNAs lacking a termination codon. Science 295: 2262–2264.

Wilson MA, Meaux S, van Hoof A. 2007. A genomic screen in yeast reveals novel aspects of nonstop mRNA metabolism. Genetics 177: 773–784.

Wu W, Liu S, Ruwe H, Zhang D, Melonek J, Zhu Y, Hu X, Gusewski S, Yin P, Small ID, et al. 2016. SOT1, a pentatricopeptide repeat protein with a small MutS-related domain, is required for correct processing of plastid 23S-4.5S rRNA precursors in Arabidopsis thaliana. The Plant Journal 85: 607–621. http://dx.doi.org/10.1111/tpj.13126.

Yang J-Y, Deng X-Y, Li Y-S, Ma X-C, Feng J-X, Yu B, Chen Y, Luo Y-L, Wang X, Chen M-L, et al. 2018. Structure of Schlafen13 reveals a new class of tRNA/rRNA-targeting RNase engaged in translational control. Nat Commun 9: 1165.

Zhang D, Burroughs AM, Vidal ND, Iyer LM, Aravind L. 2016. Transposons to toxins: the provenance, architecture and diversification of a widespread class of eukaryotic effectors. Nucleic Acids Res 44: 3513–3533.

Zhou W, Lu Q, Li Q, Wang L, Ding S, Zhang A, Wen X, Zhang L, Lu C. 2017. PPR-SMR protein SOT1 has RNA endonuclease activity. Proc Natl Acad Sci U S A 114: E1554–E1563.

Zimmermann L, Stephens A, Nam S-Z, Rau D, Kübler J, Lozajic M, Gabler F, Söding J, Lupas AN, Alva V. 2018. A Completely Reimplemented MPI Bioinformatics Toolkit with a New HHpred Server at its Core. J Mol Biol 430: 2237–2243.

